# Genetically diverse Mycobacterium tuberculosis isolates manipulate inflammasome activation and IL-1β secretion independently of macrophage metabolic rewiring

**DOI:** 10.1101/2024.06.10.598180

**Authors:** Ana Isabel Fernandes, Alexandre Jorge Pinto, Diogo Silvério, Ulrike Zedler, Carolina Ferreira, Iola F. Duarte, Ricardo Silvestre, Anca Dorhoi, Margarida Saraiva

**Affiliations:** Instituto de Investigação e Inovação em Saúde (i3S), Universidade do Porto, Porto, Portugal; Doctoral Program in Molecular and Cell Biology, Instituto de Ciências Biomédicas Abel Salazar (ICBAS), Universidade do Porto, Porto, Portugal; Institute of Immunology, Friedrich-Loeffler-Institut (FLI), Greifswald-Insel Riems, Germany; Life and Health Sciences, Research Institute (ICVS), School of Medicine, University of Minho, Braga, Portugal; ICVS/3B’s-PT Government Associate Laboratory, Braga, Portugal; CICECO – Aveiro Institute of Material, Department of Chemistry, University of Aveiro, Aveiro, Portugal; Faculty of Mathematics and Natural Sciences, University of Greifswald, Greifswald, Germany; IBMC – Instituto de Biologia Molecular e Celular, University of Porto, Porto, Portugal

## Abstract

The natural diversity of *Mycobacterium tuberculosis* is gaining relevance in dictating the outcome of tuberculosis (TB). We previously revealed a link between TB severity and *M. tuberculosis*-driven evasion of the macrophage cytosolic surveillance systems, with isolates from severe TB cases reducing inflammasome activation and interleukin (IL)-1β production by infected cells. IL-1β production and inflammasome activation are commonly associated with the metabolic reprogramming of stimulated macrophages. Thus, we questioned whether the differential modulation of the inflammasome and IL-1β by *M. tuberculosis* isolates depended on distinct macrophage metabolic reprogramming. Using metabolic inhibitors, mice deficient for key metabolic regulators, and a metabolomics approach, we found that the macrophage metabolic landscape was similar regardless of the infecting *M. tuberculosis* isolate. Paralleling single-TLR activated macrophages, inhibition of glycolysis during infection impaired IL-1β secretion. However, departing from TLR based models, in *M. tuberculosis*-infected macrophages IL-1β secretion was independent of macrophage mitochondrial metabolic changes and the transcription factor hypoxia-inducible factor (HIF)-1α. Additionally, we found a previously unappreciated impact of host metabolic inhibitors on the pathogen, and show that inhibition of the mycobacteria metabolism dampened both inflammasome activation and IL-1β production. Collectively, our study raises awareness of the potential confounding effect of host metabolic inhibitors acting on the pathogen itself and demonstrates that the modulation of the inflammasome by *M. tuberculosis* may be uncoupled from the host metabolic reprogramming.

**Author Summary:** *Mycobacterium tuberculosis* is the causative agent of tuberculosis and one of the top infectious killers in the world, with around 1.3 million deaths reported annually. The genetic variability of this pathogen can shape its interaction with the host and modulate disease outcomes. We previously found that *M. tuberculosis* clinical isolates from patients with severe forms of tuberculosis evade cytosolic surveillance systems in macrophages. Here, we explored whether this evasion tactic was linked to metabolic alterations in the infected macrophages. We found that different *M. tuberculosis* isolates induced similar metabolic changes in infected macrophages. Additionally, we demonstrate that both host glycolysis and pathogen’s metabolism were pivotal for maximum IL-1β production. These findings highlight the complexity of macrophage-pathogen interactions and emphasize that bacterial metabolism should be considered in metabolic studies and may be amenable to therapeutic intervention against tuberculosis.

## Introduction

Cellular activation and signalling cascades leading to cytokine secretion are pivotal aspects of an efficient immune response, with metabolic reprogramming of immune cells being central to this process (1–3). There has been a growing interest in understanding how the metabolic reprogramming occurs in the context of infection, namely during tuberculosis (TB), a disease that claims over 1.3 million lives annually (4).

Data from experimental mouse infections show that *Mycobacterium tuberculosis* shifts the lung immune cell metabolism from oxidative phosphorylation towards glycolysis, akin to the Warburg effect (5). In particular, the downregulation of tricarboxylic acid (TCA) cycle and electron transport chain (ETC) enzymes, alongside upregulation of glycolytic enzymes and enhanced glucose uptake, facilitates ATP generation and metabolite accumulation for anti-microbial responses (5). This metabolic shift is however not ubiquitous amongst immune cells (6). Although during *M. tuberculosis* experimental infections monocyte-derived macrophages up-regulate glycolysis, alveolar macrophages rely on β-oxidation and fatty acid uptake (5,7,8). Several in vitro studies further support the metabolic reprogramming of *M. tuberculosis-* infected macrophages (5,9–11). One aspect that remains poorly addressed concerns the impact of the natural diversity of the pathogen on the metabolic rewiring of the infected cell. Multi-drug resistant (MDR) *M. tuberculosis* strains differentially modulate the macrophage metabolism in an interferon (IFN)-β-dependent manner (12). Moreover, *M. tuberculosis* isolates with varying degrees of virulence also distinctly modulate glucose uptake to regulate macrophage viability (13). Additionally, infection of macrophages with live versus irradiated/heat-killed *M. tuberculosis* leads to distinct metabolic changes in infected cells (14–16). Collectively, these studies suggest that a link between *M. tuberculosis* virulence and the metabolic adaptation of the infected macrophage might be in place. Furthermore, a crosstalk between NLRP3 activation, interleukin (IL)-1β production, and the macrophage’s metabolic reprogramming has been established. Alterations in inflammasome activation have been linked to the accumulation of TCA intermediates, such as succinate and itaconate (17,18). The recognition of live bacteria by macrophages through NLRP3-dependent pathways transiently decreases ETC complexes assembly and induces mitochondrial ROS production (19). In turn, inhibition of ETC complexes prevents NLRP3 activation, in a phosphocreatine-dependent manner (20).

Previous work from our group revealed that *M. tuberculosis* clinical isolates recovered from patients with severe forms of TB, evaded cytosolic surveillance systems in macrophages, resulting in lower inflammasome activation and IL-1β production. In contrast, clinical isolates retrieved from patients with mild TB disease activated the inflammasome in macrophages, thus inducing high levels of IL-1β (21). In this study, we investigated whether the differential NLRP3 activation and IL-1β production induced in macrophages by genetically diverse *M. tuberculosis* resulted from distinct host metabolic reprogramming. We found that, regardless of their ability to activate the inflammasome and induce IL-1β, the tested *M. tuberculosis* isolates triggered a similar metabolic shift in infected macrophages. In essence, activation of the glycolytic pathway was pivotal for maximum IL-1β production, but in contrast to what is described for single TLR agonists (17,22,23), in live *M. tuberculosis* infections, mitochondria metabolic alterations were uncoupled from the inflammasome activation and did not impact IL-1β secretion by infected cells. Furthermore, blockade of the pathogen metabolism using host-related or pathogen-specific inhibitors abrogated the inflammasome activation and subsequent IL-1β production. Collectively, our study highlights the importance of studying metabolic/inflammatory networks using complex agents and raises awareness of the potential impact of host metabolic inhibitors on the pathogen.

## Results

### Genetically distinct *M. tuberculosis* clinical isolates elicit similar glycolytic reprogramming in macrophages

We have previously reported a bacteria-driven link between the severity of TB and IL-1β production by infected macrophages and have identified two *M. tuberculosis* isolates, 4I2 (high IL-1β inducer; mild TB) and 6C4 (low IL-1β inducer; severe TB), as prototypes of such differences (Fig 1A) (21). Given the link between macrophage metabolic reprogramming and inflammasome activation (2,3,22), we questioned whether these genetically diverse clinical isolates of *M. tuberculosis* may also differentially control the macrophage metabolic shift with an impact on IL-1β release.

**Fig 1.**
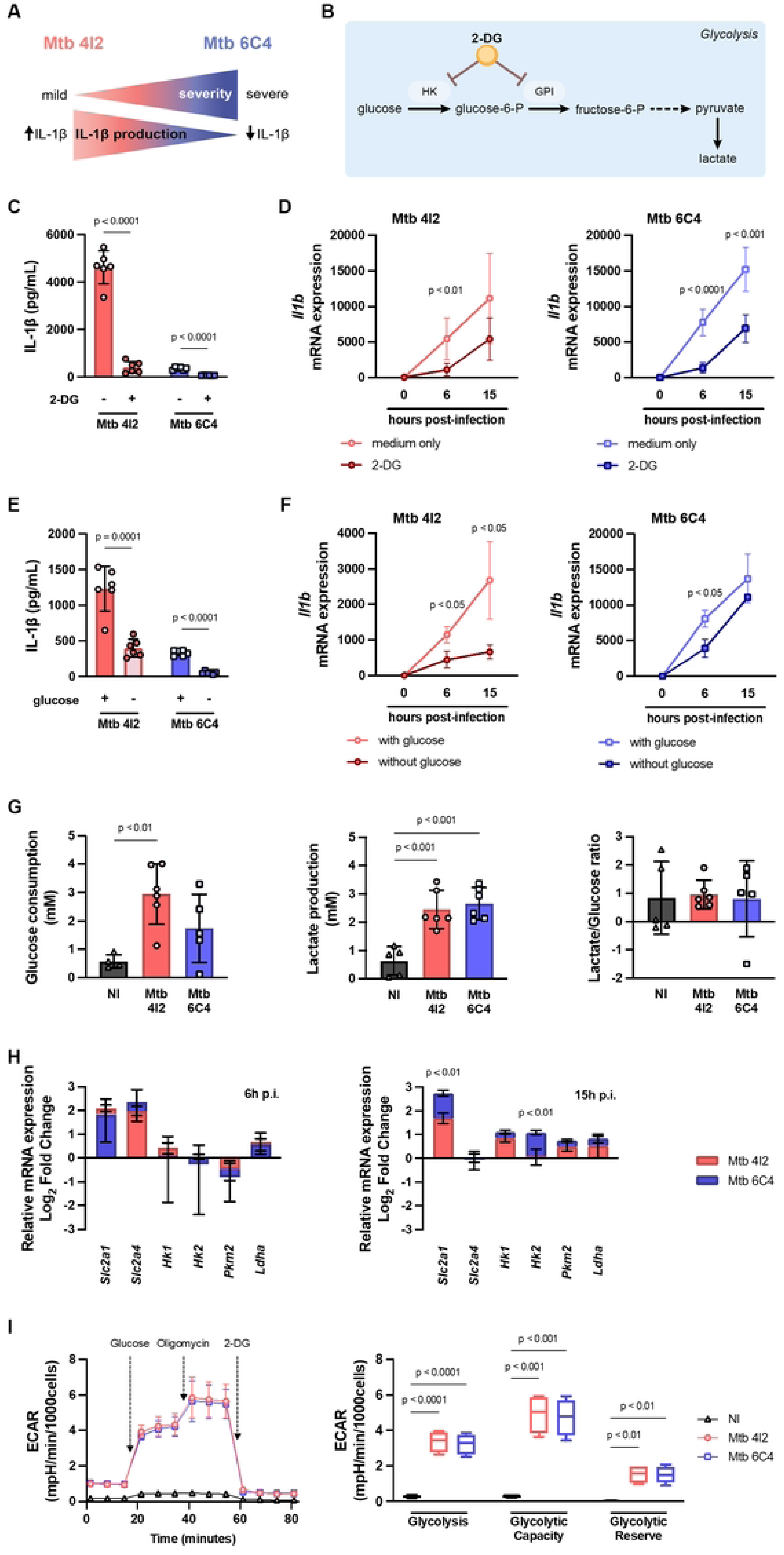
Glucose consumption contributes to IL-1β production by infected macrophages but glycolysis does not differ between the two *M. tuberculosis* clinical isolates. (**A**) Inverse correlation between disease severity and IL-1β production in cells infected with *M. tuberculosis* 4I2 and *M. tuberculosis* 6C4, as described in (21). (**B**) 2-DG inhibitory action within the glycolytic pathway. (**C-F**) Mouse BMDM were left untreated (−) or pre-treated (+) for 2h with 2-DG (**C, D**) or incubated with DMEM without glucose (**E, F**) and subsequently infected at an MOI of 2 with either clinical isolate. (**C, E**) The amount of secreted IL-1β was quantified in the cell culture supernatants 24h post-infection. (**D, F**) 6h and 15h post-infection, cell cultures were lysed and the levels of *Il1b* determined by real-time PCR. (**G-H**) BMDM from C57BL/6 mice were infected at an MOI of 2 with either clinical isolate. (**G**) Glucose consumption and lactate production were measured through HPLC in the supernatant of infected cultures after 24h, and the lactate/glucose ratio was calculated based on these values. (**H**) 6h and 15h post-infection, cell cultures were lysed and the expression of the indicated genes determined by real-time PCR. Data are represented in log_2_(fold change), with 0 representing non-infected macrophages. (**I**) BMDM infected at MOI of 2 for 24h were submitted to glycolysis stress test and ECAR was measured with an XFp Analyser. Glycolysis, glycolytic capacity, and glycolytic reserve were calculated. Represented is the mean ± SD of triplicate (**C-H**) or duplicate (**I**) wells for two experiments (**C-E, G, I**) or one experiment representative of two (**F, H**). For ECAR assays (**I**) data are also represented in a box and whiskers plot (Tukey’s method). Statistical analysis was performed using two-tailed unpaired Student’s *t*-test (**C-F, H**) or One-way ANOVA (**G, I**). 2-DG – 2-deoxy-glucose, ECAR – extracellular acidification rate, GPI – glucose-6-phosphate isomerase, HK – hexokinase, Mtb – *M. tuberculosis*, NI – non-infected, p.i. – post-infection.

We first evaluated whether glycolysis affected IL-1β production by bone-marrow-derived macrophages (BMDM) infected with either clinical isolate. As we reported (21), the differences in IL-1β production by macrophages infected with *M. tuberculosis* 4I2 or *M. tuberculosis* 6C4 were due to post-transcriptional alterations (S1A Fig) and dependent on inflammasome activation, as shown in the presence of MCC950, an NLRP3 inhibitor (S1B Fig). Inhibition of glycolysis with 2-DG, a non-metabolizable glucose analogue (Fig 1B), abrogated IL-1β secretion by BMDM infected with either clinical isolate (Fig 1C). The impact of 2-DG was also observed at the transcriptional level, as infected BMDM expressed lower levels of *Il1b* in the presence of the inhibitor, irrespective of the *M. tuberculosis* isolate (Fig 1D). In line with these data, secretion of IL-1β (Fig 1E) and transcription of the *Il1b* gene (Fig 1F) were also decreased in *M. tuberculosis* 4I2- or 6C4-infected BMDM cultures deprived of glucose. Moreover, BMDM infected in the presence of 2-DG or in the absence of glucose showed a significant decrease in lactate, a by-product of glycolysis (S1C Fig). Of note, downstream inhibition of the pentose phosphate pathway with 6-animonicotinamide (S1D Fig) did not significantly impact IL-1β secretion by infected BMDM (S1E Fig). Importantly, none of the used inhibitors altered cell viability (S1F Fig).

Given that BMDM infected with *M. tuberculosis* 4I2 or 6C4 required glycolysis to produce IL-1β, we next questioned if the macrophage glycolytic shift induced by either isolate might be different, thus providing an explanation for the differential induction of IL-1β. When compared to non-infected cells, *M. tuberculosis* infected BMDM showed increased glucose consumption and lactate production, with similar lactate/glucose ratios as measured by HPLC (Fig 1G). Consistently, the expression of genes encoding glucose transporters and glycolytic enzymes measured by real-time PCR was increased over time for both infections, with little to no differences between them (Fig 1H). The only significant differences were observed for the expression of *Slc2a1* and *Hk2* which were higher in infections with *M. tuberculosis* 6C4 at a later time point (15 hours post-infection; Fig.1H). Furthermore, variations in several glycolytic parameters of BMDM were measured in real-time upon infection with either live *M. tuberculosis* clinical isolate, using the extracellular acidification rate (ECAR) assay. In line with previous results, we observed increased glycolysis, glycolytic capacity, and glycolytic reserve in infected BMDM, all of which were comparable between *M. tuberculosis* 4I2 and 6C4 (Fig 1I). Altogether, our data indicate a common glycolytic response to live *M. tuberculosis* infection regardless of downstream strain-specific differences in inflammasome activation and IL-1β production.

### IL-1β production by infected macrophages is independent of mitochondrial ROS and mTOR

Given that the differences in secretion of IL-1β by macrophages infected with *M. tuberculosis* 4I2 or 6C4 was not related to differential glucose uptake or glycolytic shift, we next investigated the potential contribution of other metabolic pathways. The accumulation of TCA intermediates, such as succinate, in macrophages upon bacterial infection has been reported (24–26). Succinate oxidation may lead to mitochondrial depolarization, mitochondrial ROS production and activation of the mTOR pathway, all converging to inflammasome activation (23,27) (Fig 2A). To assess the impact of mitochondrial metabolic alterations in IL-1β production by macrophages infected with *M. tuberculosis* 4I2 or 6C4, we inhibited the enzyme succinate dehydrogenase (SDH) with dimethyl malonate (DMM; Fig 2A). We observed a marked decrease in IL-1β protein in DMM-treated BMDM irrespective of the *M. tuberculosis* clinical isolate (Fig 2B). However, in contrast to what was observed when blocking glycolysis (Fig 1), inhibition of SDH did not impact *Il1b* transcription levels (Fig 2C). This finding suggests a post-transcriptional contribution of SDH activity for IL-1β production, compatible with NLRP3 activation linked to the mitochondrial respiratory chain (Fig 2A), which we investigated next. Surprisingly, staining of macrophages with Mitotracker Red CMXRos revealed no differences in mitochondrial membrane potential between non-infected and infected BMDM, and between the two *M. tuberculosis* isolates (S2A Fig), suggesting that mitochondria remained polarized upon infection. Additionally, infection of BMDM with either clinical isolate downregulated genes encoding enzymes from the ETC (S2B Fig). This is in line with other reports that live bacteria, but not LPS, transiently impacts ETC enzymes (19). Functional sequestration of mitochondrial ROS with mitoTEMPO did not impact IL-1β secretion (Fig 2D) or transcription (Fig 2E). Likewise, blockade of the mTOR pathway with rapamycin resulted in unaltered IL-1β secretion (Fig 2F) or transcription (Fig 2G). As seen with glycolytic inhibitors, none of the used mitochondrial inhibitors altered cell viability (S2C Fig).

**Fig 2.**
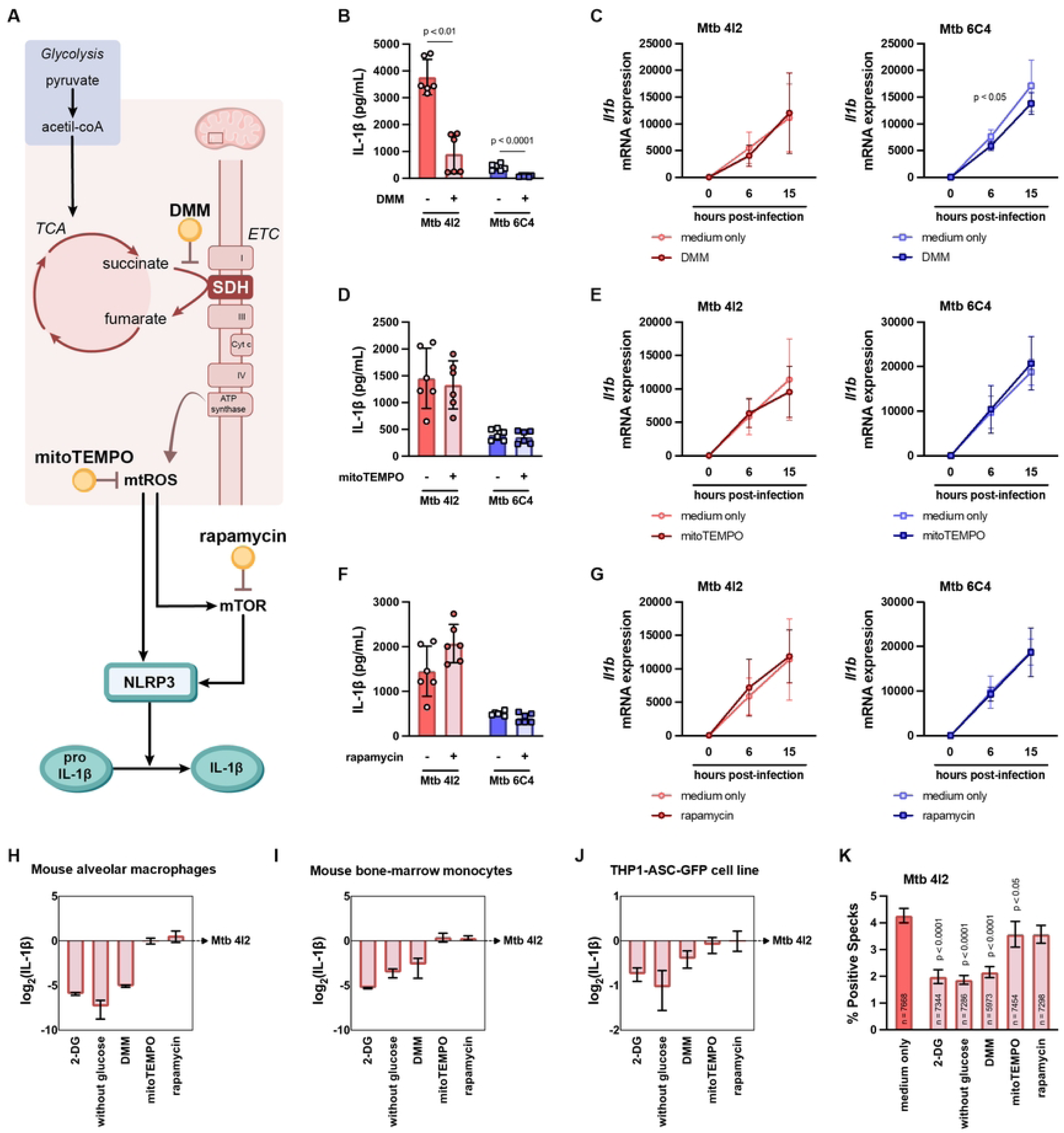
Dimethyl malonate inhibits inflammasome activation and IL-1β production independently of mitochondrial ROS and mTOR. **(A)** DMM, mitoTEMPO and rapamycin inhibitory action within the metabolic pathway. (**B-G**) BMDM from C57BL/6 mice were left untreated (−) or pre-treated (+) for 2h with DMM (**B, C**), mitoTEMPO (**D, E**), or rapamycin (**F, G**) and subsequently infected with either clinical isolate at an MOI of 2. (**B, D, F**) The amount of secreted IL-1β was quantified in the cell culture supernatants 24h post-infection. (**C, E, G**) 6h and 15h post-infection, cell cultures were lysed and *Il1b* expression measured by real-time PCR. (**H**) Alveolar macrophages, (**I**) bone-marrow-derived monocytes from C57BL/6 mice, and (**J**) THP1-ASC-GFP macrophages, were left untreated (−) or pre-treated (+) with the indicated inhibitors for 2h and infected with *M. tuberculosis* 4I2 at an MOI of 1 for 24h. The amount of secreted IL-1β was quantified in the cell culture supernatants. Data are represented in log_2_(fold change), with 0 representing macrophages infected with *M. tuberculosis* 4I2. (**K**) PMA-differentiated THP-1-ASC-GFP were treated with inhibitors for 2h as indicated in the figure and infected with *M. tuberculosis* 4I2 at an MOI of 1 and the percentage of speck-positive cells determined 6h post-infection. Represented is the mean ± SD of triplicate wells for two experiments (**B-G, I, J**) or one experiment (**H, K**). For immunofluorescence, 60 fields per replicate (total number of cells indicated in each bar) were acquired and analysed. Statistical analysis was performed using two-tailed unpaired Student’s *t*-test (**B-G**) or One-way ANOVA (**K**). 2-DG – 2-deoxy-glucose, DMM – dimethyl malonate, ETC – electron transport chain, Mtb – *M. tuberculosis*, mTOR – mammalian target of rapamycin, mtROS – mitochondrial ROS, SDH – succinate dehydrogenase, TCA – tricarboxylic acid.

These findings were surprising as they suggest that IL-1β secretion by *M. tuberculosis-*infected BMDM does not depend on alterations to the ETC, unlike what has been reported in other systems based on single TLR macrophage activation (20,22). To investigate whether these results extended beyond BMDM, mouse alveolar macrophages (Fig 2H) or bone-marrow monocytes (Fig 2I), and macrophages differentiated from the human monocytic cell line THP1 (Fig 2J) were infected with *M. tuberculosis* 4I2 in medium alone or in the presence of the aforementioned inhibitors. As observed for BMDM, only the inhibition of glycolysis (with 2-DG or in medium without glucose) or of SDH (with DMM) inhibited IL-1β secretion (Fig 2H-2J). Consistent with this, a decrease in inflammasome activation was detected in infected THP1-ASC-GFP reporter cells in the case of glycolytic (2-DG and medium without glucose) or SDH (DMM) inhibition (Fig 2K), whereas mitochondrial ROS sequestration and mTOR inhibition resulted in a sustained percentage of ASC-speck-positive cells (Fig 2K). Similarly to our previously published data (21), the percentage of ASC-speck-positive cells was higher in the case of *M. tuberculosis* 4I2 infections when compared to *M. tuberculosis* 6C4 or non-infected cells (S2D Fig) and direct inflammasome inhibition with MCC950 reduced the percentage of positive specks detected (S2D Fig).

Collectively, our findings suggest that despite the SDH contribution to IL-1β secretion, the canonical pathways linking mitochondrial metabolism and inflammasome activation upon single TLR stimulation are likely to operate differently upon complex stimuli, such as live *M. tuberculosis*.

### *M. tuberculosis* clinical isolates elicit similar metabolomic profiles in infected macrophages

To gain further insights into possible metabolic mediators underlying why SDH inhibition decreased the secretion of IL-1β by infected macrophages, we next performed an NMR metabolomic analysis of the cell culture supernatants (extracellular metabolome) and their corresponding cell extracts (intracellular metabolome). We analysed BMDM cultures (non-infected, infected with *M. tuberculosis* 4I2 or with *M. tuberculosis* 6C4), as well as cultures of the clinical isolates obatined under the same conditions but in the absence of host cells. We also included in our analysis a condition of medium alone, which was used to define the basal levels of the different metabolites, with values below or above this baseline representing consumption or excretion, respectively.

Principal component analysis (PCA) scores of the extracellular metabolome showed the infected cells clustering closely together (Fig 3A), suggesting that the two *M. tuberculosis* isolates under study are likely to reprogram the macrophage metabolism in similar ways. Consistent with our previous results, *M. tuberculosis*-infected BMDM consumed more glucose and secreted more lactate than non-infected macrophages, irrespective of the infecting clinical isolate (Fig 3B). This was accompanied by a slight increase in pyruvate consumption and no effect on fructose, two other glycolytic metabolites detected (Fig 3B). As for TCA metabolites, an increase of released itaconate and glutamate was detected in infected BMDM cultures, again with no differences between *M. tuberculosis* isolates (Fig 3C). As for the supernatants obtained upon bacteria cultured in the absence of BMDM, the vast majority of the detected metabolites was similar between the two isolates and mostly unaffected as compared to the culture medium (Fig 3B-3C and S3 Fig). The only exception to this were pyruvate and asparagine, which were more consumed by *M. tuberculosis* 6C4-infected cells (S3A Fig).

**Fig 3.**
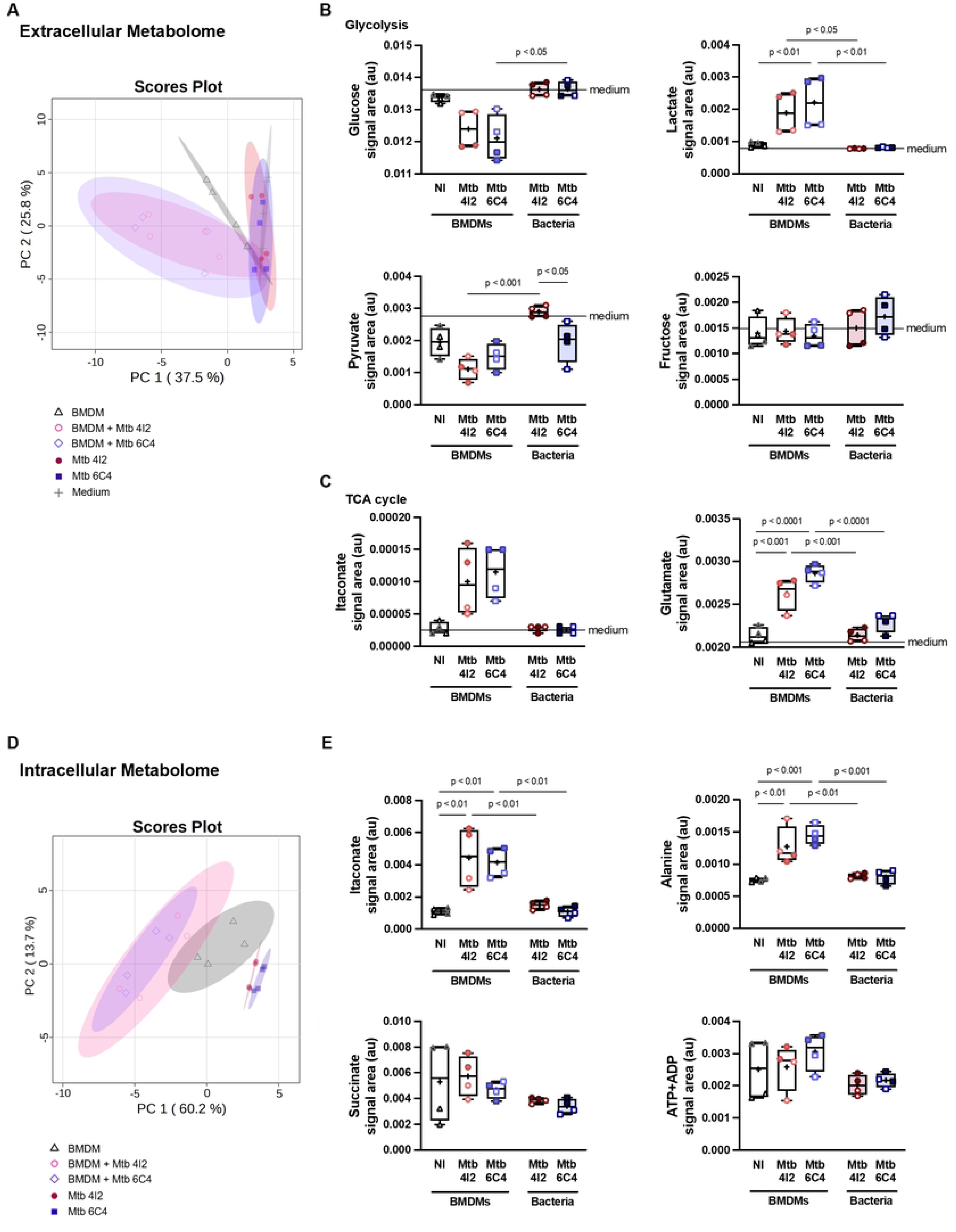
NMR-based metabolomic analysis of macrophages infected with *M. tuberculosis* clinical isolate 4I2 or 6C4 reveals similar metabolic profiles. (**A**) Principal component analysis (PCA) of spectral data from the culture supernatants of non-infected BMDM, macrophages infected with *M. tuberculosis* 4I2 or 6C4 for 24h at an MOI of 2, and *M. tuberculosis* 4I2 or 6C4 cultured in similar medium but in the absence of host cells. (**B-C**) Representative metabolites consumed and excreted among the tested conditions. Variations are expressed in relation to media only. (**D**) Principal component analysis (PCA) of spectral data recorded for intracellular aqueous extracts the same conditions as in (**A**). (**E**) Relative intracellular levels of representative metabolites across the tested conditions. Represented are box and whiskers plots (Tukey’s method) of duplicate wells for two experiments. Each replicate is the pool of 24 wells in a 24 well-plate. Statistical analysis was performed using One-way ANOVA. au – arbitrary unit, BMDM – bone-marrow derived macrophages, Mtb – *M. tuberculosis*, NI – non-infected, TCA - tricarboxylic acid.

As for the analysis of the intracellular metabolome, the PCA plot showed some separation between cellular and bacteria-only conditions (Fig 3D). However, infected cells clustered together irrespective of the infecting *M. tuberculosis* clinical isolate, supporting that no substantial differences exist between these two conditions. In line with the extracellular metabolome data and with what has been described for other bacterial infections (28–30), we detected an increase of itaconate and alanine in infected cells which was similar for *M. tuberculosis* 4I2 and 6C4 infections (Fig 3E). Additionally, we did not detect an accumulation of intracellular succinate or ATP+ADP upon infection (Fig 3E). Of note, we also detected a series of other metabolites some of which were modulated by the infection, but always in a similar manner independently of the *M. tuberculosis* isolate (S3 Fig).

In essence, for both *M. tuberculosis* 4I2 and 6C4 infections of BMDM, we observed similar metabolic alterations, including a similar glycolytic shift and increase in by-products of the TCA cycle which did not include succinate.

### IL-1β production by *M. tuberculosis* infected macrophages is independent of HIF-1α and IRG1 activity

Succinate has been described to stabilize hypoxia-inducible factor (HIF)-1α, a transcription factor for *Il1b* (17,31) whereas itaconate has been associated with the inhibition of SDH and NLRP3 inflammasome (18,32) (Fig 4A). We thus tested the contribution of these two pathways for IL-1β production by macrophages upon *M. tuberculosis* infection. Infection of BMDM generated from myeloid-restricted HIF-1α deficient (mHIF1α^-/-^) mice (33) with either *M. tuberculosis* isolate yielded no effect on IL-1β protein levels when compared to WT mice (Fig 4B), with only minor differences seen in the amounts of *Il1b* mRNA (Fig 4C). To test a possible role for itaconate, we generated BMDM from IRG1^-/-^ mice (34), which fail to convert aconitate into itaconate. Lack of IRG1 did not impact IL-1β secretion (Fig 4D) or transcription (Fig 4E) by BMDM infected with either clinical isolate. In conclusion, all our findings demonstrate a similar metabolic reprogramming of the macrophage in response to two distinct *M. tuberculosis* isolates, and, except for the effect of DMM, point to a minor contribution of the TCA cycle to IL-1β production by infected macrophages. Our data thus highlight that differential inflammasome activation and IL-1β production may, at least in certain systems, be independent of the macrophage mitochondria metabolic rewiring.

**Fig 4.**
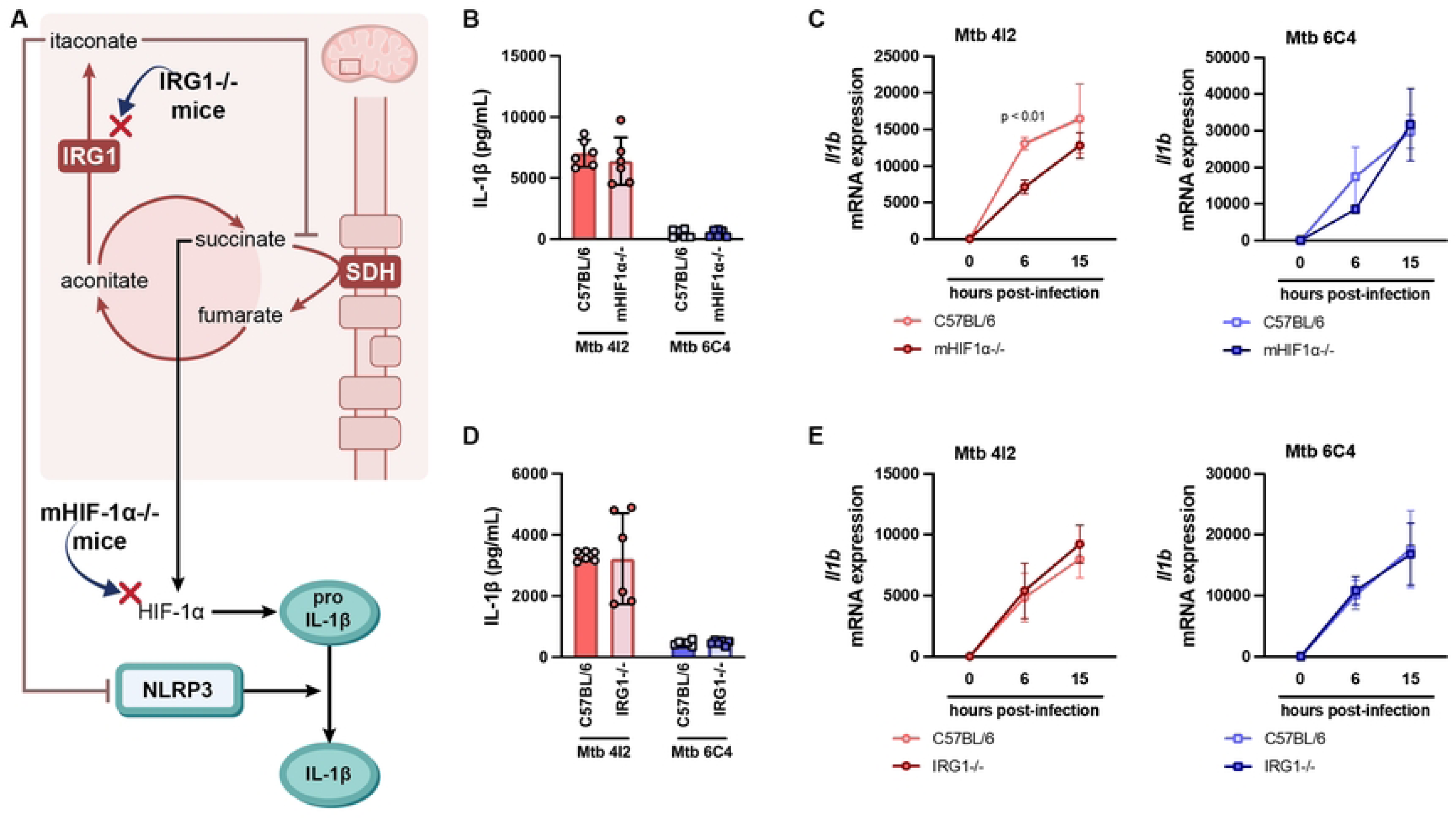
IL-1β production by *M. tuberculosis* infected macrophages is independent of HIF-1α and IRG1 activity. (**A**) HIF-1α and IRG1 pathways and their impact on IL-1β regulation. BMDM from C57BL/6 and mHIF-1α^-/-^ (**B, C**) or IRG1^-/-^ (**D, E**) mice were infected with either clinical isolate at an MOI of 2 for 24h. (**B, D**) The amount of secreted IL-1β was quantified in the cell culture supernatants 24h post-infection. (**C, E**) 6h and 15h post-infection, cell cultures were lysed and the *Il1b* expression measured by real-time PCR. Represented is the mean ± SD of triplicate wells for two experiments (**B, D, E**) or one experiment (**C**) representative of two. Statistical analysis was performed using a two-tailed unpaired Student’s *t*-test. HIF-1α – hypoxia inducible factor 1 subunit alpha, IRG1 – immune responsive gene 1, Mtb – *M. tuberculosis*, SDH – succinate dehydrogenase.

### DMM acts on the pathogen to inhibit IL-1β secretion by infected macrophages

We next investigated in more detail how DMM might interfere with IL-1β production by infected cells, since our described findings did not support a role for the host metabolic pathways in this process. The effect of SDH inhibition by DMM was post-transcriptional, not related to the usual host metabolic molecular players, and involved inflammasome inhibition (Fig 2). Moreover, it paralleled the effect observed when the bacterial metabolism was inhibited by rifampicin during infection (21). Taking all these results into consideration, we hypothesised that DMM might primarily interfere with the bacteria, not the host, metabolism.

*M. tuberculosis* encodes three different SDH enzymes, all predicted to be functionally capable of oxidising succinate. These are SDH-1 (Rv0247c to Rv0249c), SDH-2 (Rv3316 to Rv3319), and fumarate reductase (Rv1552 to Rv1555) (35–37). We focused on SDH-1 due to its high homology to the *Mycobacterium smegmatis-*encoded molecule, which has a published structure and predicted electron transfer mechanism (38). We predicted the structure of *M. tuberculosis* SDH-1 using ColabFold (39) and superpositioned it with the resolved structure of *M. smegmatis* SDH-1 containing ubiquinone-1 (instead of menaquinone) as the ligand (PDB: 7D6V) (Fig 5Ai). The electron transfer pathway in *M. smegmatis* SDH-1 is predicted to occur from the oxidation of succinate in the flavoprotein domain (represented in cyan, Fig 5Ai and ii), to the quinone binding site (represented in green, Fig 5Ai and ii), by a chain of redox centers including the flavin-adenine nucleotide (FAD), the [2Fe-2S], [4Fe-4S], [3Fe-4S] iron-sulphur cluster, and the Rieske-type [2Fe-2S] cluster (represented in yellow, Fig 5Ai and ii) (38). Given the structural similarities between the *M. smegmatis* and *M. tuberculosis* molecules, their electron transfer mechanism is likely similar. Furthermore, due to the overall arrangement of redox centers, they may function akin to the mammalian SDH (40), and could be inhibited by the same small molecules, thus making the bacterial SDH a possible target for the inhibitor DMM. The mode of action of DMM remains unclear, but based on the analysis of the structure superposition, it likely involves direct binding to the quinone-binding domain of SDH in place of ubiquinone-1 (Fig 5Aii and iii), or acting as a pro-drug which is converted into malonate and competes for the succinate binding site in SDH (Fig 5Aii and iii) (41).

**Fig 5.**
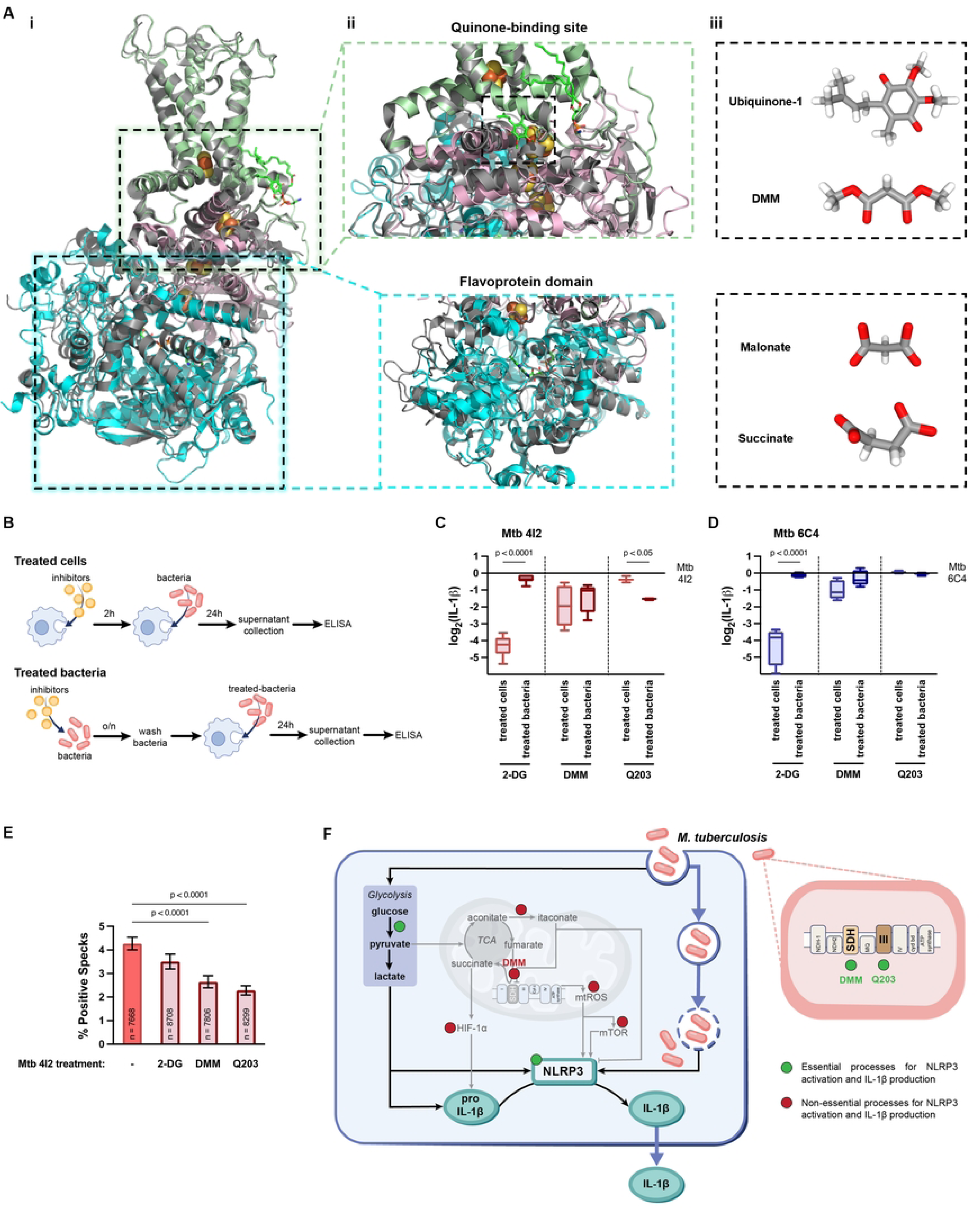
Infection of macrophages with *M. tuberculosis* 4I2 exposed to dimethyl malonate results in reduced inflammasome activation and IL-1β production. (**A**) *M. tuberculosis* SDH-1 protein structure predicted with ColabFold superpositioned with *M. smegmatis* SDH-1. Flavoprotein domain (represented in cyan), Iron-sulphur domain (represented in pink) and Membrane anchor domain (represented in green) are superpositioned with SDH-1 from *M. smegmatis* in complex with ubiquinone-1 (7D6V, grey). Ubiquinone-1 (CID 4462), DMM (CID 7943), malonate (CID 9084) and succinate (CID 160419) structures were extracted from PubChem. Amino acid sequences for *M. tuberculosis* H37Rv used in ColabFold were extracted from the NCBI protein database: Flavoprotein subunit (NP_214762.1); Iron-Sulphur subunit (NP_217836.1); Membrane anchor subunit (NP_214763.1). (**B**) Schematic representation of the protocol for treating bacteria or macrophages with used metabolic inhibitors. *M. tuberculosis* 4I2 (**C**) and 6C4 (**D**) or C57BL/6 BMDM were pre-treated with the indicated inhibitors according to the protocol in **B.** Treated macrophages were infected with non-treated bacteria and non-treated macrophages were infected with treated bacteria, at an MOI of 2 for 24h. The amount of secreted IL-1β was quantified in the cell culture supernatants. Data are represented in log_2_(fold change), with 0 representing non-treated macrophages infected with non-treated *M. tuberculosis* 4I2 (**C**) or 6C4 (**D**). (**E**) THP1-ASC-GFP macrophages were infected with treated *M. tuberculosis* 4I2, at an MOI of 1 for 6h and the percentage of speck-positive cells was determined for the different conditions. Represented are box and whiskers plots (Tukey’s method) (**C, D**) or the mean ± SD (**E**) of triplicate wells for two experiments (**C, D**) or one experiment (**E**). For immunofluorescence, 60 fields (total number of cells indicated in each bar) per replicate were acquired and analysed. Statistical analysis was performed using One-way ANOVA (**H**). (**F**) Model of the macrophage immunometabolic adaptation during *M. tuberculosis* infection and its interaction with the inflammasome activation and IL-1β secretion. 2-DG – 2-deoxy-glucose, DMM – dimethyl malonate, HIF-1α – hypoxia inducible factor 1 subunit alpha, Mtb – *M. tuberculosis*, mTOR – mammalian target of rapamycin, mtROS – mitochondrial ROS, o/n – overnight, TCA – tricarboxylic acid.

To distinguish the impact of DMM on the host versus bacteria, we compared the effect of exposing the macrophages to DMM during infection (which would affect the host, and potentially the pathogen) versus that of treating the pathogen with DMM and washing out the inhibitor prior to the infection (thus mainly targeting the pathogen; Fig 5B). We additionally included the glycolytic inhibitor 2-DG and Q203, an *M. tuberculosis*-specific inhibitor of the respiratory chain (42). As before (Fig 1), exposure of the infected cultures to 2-DG highly impacted the secretion of IL-1β by BMDM independently of the *M. tuberculosis* isolated used (Fig 5C-5D). However, pre-treating *M. tuberculosis* with 2-DG did not impact IL-1β levels showing that the glycolysis requirement for IL-1β production results from the effect of 2-DG in the host (Fig 5C-5D). In contrast, the impact of DMM on IL-1β secretion was similar for the two experimental conditions (Fig 5C-5D) suggesting that affecting the bacteria was enough to decrease IL-1β secretion. Moreover, this effect of DMM in *M. tuberculosis* was higher in the case of isolate 4I2 than of isolate 6C4, fitting with our previous data suggesting that the activation of the inflammasome by *M. tuberculosis* 4I2 requires metabolically active bacteria (21). Finally, Q203 only impacted pathogen-mediated IL-1β secretion (Fig 5C), again in line with our previous findings using rifampicin (21). Of note, BMDM infected with DMM-treated bacteria showed no differences in *Il1b* transcript when compared to DMM-treated BMDM (S4 Fig), further supporting its post-transcriptional effect. Finally, we measured the inflammasome activation in THP1-ASC-GFP reporter cells infected with *M. tuberculosis* 4I2 pre-treated or not with each of these inhibitors. Treatment of *M. tuberculosis* 4I2 with DMM or Q203 decreased the activation of the host inflammasome, with no significant effect detected when the pathogen was treated with 2-DG (Fig 5E). Collectively, these data suggest that the effect of DMM is likely mediated by inhibition of the pathogen metabolism, specifically by interfering with the ability of certain isolates of *M. tuberculosis* to activate the host inflammasome and subsequent IL-1β production by infected macrophages. In summary (Fig 5F), our results position the macrophage glycolytic shift promoted by *M. tuberculosis* infection and the bacteria metabolism as critical steps for IL-1β production, with no apparent contribution of the host TCA cycle and ETC, contrasting previously described findings for single TLR agonists.

## Discussion

Recent research has highlighted the significant impact of *M. tuberculosis* diversity on host-pathogen interactions and TB outcomes. Despite highly similar, closely related *M. tuberculosis* strains employ distinct and highly effective mechanisms to subvert host immune responses. These mechanisms range from receptor recognition to inflammasome activation and type I interferon responses (12,13,21,43–45), also likely involving the metabolic reprogramming of infected cells (6). Although it is widely accepted that *M. tuberculosis* infection shifts macrophage metabolism from oxidative phosphorylation to glycolysis, how genetically diverse *M. tuberculosis* isolates may modulate this process remains largely unexplored (10,11,14,16,46). To answer this question, we used a previously established system focused on live, genetically diverse clinical isolates that distinctly modulate NLRP3 activation and IL-1β production in macrophages in a host-independent manner (21).

Here, we report that the two *M. tuberculosis* clinical isolates under study elicited similar glycolytic responses in infected macrophages, indicating a shared bioenergetic profile despite downstream strain-specific differences in inflammasome activation and IL-1β production. This suggests that while bacterial genetic variation may impact the macrophage cytokine response, the energetic requirements to mount this initial immune response are similar and conserved. Additionally, our study underpins the glycolytic pathway as a crucial regulator of IL-1β production during *M. tuberculosis* infection, impacting both *Il1b* transcription and NLRP3 inflammasome activation. This is in line with previously published data, as hexokinase, a glycolytic enzyme, has been reported to trigger the inflammasome by acting as a pattern recognition receptor (47,48).

Having established that the macrophage glycolytic shift was key for IL-1β production but not dependent on the infecting *M. tuberculosis* isolate, we next investigated a possible crosstalk between the TCA cycle and inflammasome activation. Infection of BMDM in the presence of DMM resulted in a significant decrease in IL-1β production, due to post-transcriptional modulation. This result implicated a possible contribution of the TCA remodelling to IL-1β production, as previously established mainly with LPS (17,31,32). However, it was surprising to find that sequestration of mitochondrial ROS, inactivation of the mTOR pathway, or deletion of HIF-1α did not impact IL-1β secretion by infected BMDM. These mechanisms were previously reported to link TCA cycle remodelling to inflammasome activation and maximum IL-1β production by LPS-activated macrophages (18,23,47). Departing from current immunometabolic models based on a wealth of studies performed with single TLR activation, our findings now suggest that during infection of macrophages with live *M. tuberculosis,* the inflammasome activation may be uncoupled from the TCA cycle.

Through metabolomic analyses, we found an overwhelmingly similar macrophage metabolic profile in response to the two *M. tuberculosis* under study, supporting that macrophage metabolic rewiring and differential IL-1β production are independent events. *M. tuberculosis-* infected macrophages accumulated itaconate, a metabolite involved in the regulation of succinate accumulation, mitochondrial respiration, and cytokine production with NLRP3 modulation (18,32). However, the levels of itaconate induced were similar for both *M. tuberculosis* isolates and *Irg1* abrogation did not impact IL-1β production. These findings further support the hypothesis that canonical pathways conventionally linked to inflammasome activation may operate differently under single (e.g., LPS) versus complex stimuli, as is the case of live *M. tuberculosis*.

The observation that DMM inhibited inflammasome activation and IL-1β production by *M. tuberculosis* infected macrophages without the contribution of other common host metabolites was puzzling. In our previous study, we showed that high IL-1β induction in infected macrophages depended on the ability of certain *M. tuberculosis* isolates to activate the inflammasome and demonstrated that live, metabolically active bacteria are required for this effect (21). Thus, we raised the hypothesis that DMM might be interfering with the pathogen metabolism rather than the host macrophage metabolic adaptation to infection. Combining structural prediction of the *M. tuberculosis* SDH with stimulation of macrophages with *M. tuberculosis* previously exposed to DMM, we concluded that the effect of this inhibitor on IL-1β may indeed be mediated by the pathogen. Whether DMM alters mycobacterial replication capacity, production or abundances of virulence factors, i.e. lipids (PDIM), serine/threonine kinases (PknF) or ESX-1 components, required for perturbations of the phagosomal membrane and subsequent NLRP3 activation remains to be established. Furthermore, we show that similar effects are obtained when a pathogen-specific respiratory chain inhibitor (Q203; (42)) is used. Given the essentiality of the pathogen’s metabolic activity to the activation of the inflammasome and subsequent IL-1β secretion, it is possible that the distinct phenotypes assigned to *M. tuberculosis* 4I2 and 6C4 may depend on specific metabolic characteristics of each isolate. Interesting candidate pathways to pursue are those linked to pyruvate and asparagine consumption, which we detected as differential between the two isolates.

In conclusion, our study provides valuable insights into the complex relationship between *M. tuberculosis* infection, macrophage metabolism, and IL-1β production. We demonstrate that while genetically distinct *M. tuberculosis* clinical isolates induce similar metabolic reprogramming in infected macrophages, the regulation of IL-1β production during infection relies on the macrophage glycolytic shift but is primarily governed by bacteria-specific mechanisms. It will be interesting to explore these mechanisms in the future and unveil the molecular details of this novel *M. tuberculosis* immune modulation strategy. The findings presented in this study emphasize the complexity of macrophage-*M. tuberculosis* interactions and also underscore the importance of considering bacterial metabolism as a confounding factor in these studies and as a potential target for therapeutic intervention in combating this global health threat.

## Materials and Methods

### Animals

C57BL/6 WT male or female mice between the ages of 8 to 12 weeks were used to generate BMDM, maintained and provided by the animal facilities at i3S or FLI. Animals were kept under specific-pathogen-free conditions, with controlled temperature (20–24 °C), humidity (45– 65%), light cycle (12h light/dark) and ad libitum food and water. Hif1a^fl/fl^-LysMcre^+/+^ (mHIF-1α^-/-^) and C57BL/6NJ-Acod1^em1(IMPC)J^/J (IRG1^-/-^), and matched C57BL/6 WT controls were provided by Dr Ricardo Silvestre and Dr Agostinho Carvalho, ICVS, Braga, Portugal. All experiments were performed in strict accordance with the recommendation of European Union Directive 2010/63/EU and previously approved by the Portuguese National Authority for Animal Health—*Direção Geral de Alimentação e Veterinária* (DGAV) and in accordance with current German Animal Welfare regulations. Experiments were carried out according to ARRIVE guidelines (https://arriveguidelines.org).

### Preparation of bacterial stocks

*M. tuberculosis* 4I2 and 6C4 were grown in 200 mL of Middlebrook 7H9 (BD Biosciences) liquid medium supplemented with 10% Oleic Albumin Dextrose Catalase (OADC) growth supplement and 0.2% glycerol. Bacterial suspensions were aliquoted in cryovials, frozen, and stored at –80 °C. Bacterial quantification was performed by thawing three to five vials and plating several serial dilutions in 7H11 (BD Biosciences) agar medium supplemented with 10% OADC and 0.5% glycerol. The plates were incubated for 21 days at 37 °C before colony count.

### Generation of bone-marrow-derived macrophages

Mice tibias and femurs were collected, and the bone marrow flushed with complete (c)DMEM (DMEM supplemented with 10% fetal bovine serum (FBS), 1% sodium pyruvate, 1% HEPES and 1% L-glutamine; all GIBCO). BMDM were counted and plated in Petri dishes with 100 mm diameter at the concentration of 0.5×10^6^ cells mL^−1^ in 8 mL with 20% L-929 conditioned media (LCCM) or 40 ng mL^-1^ of mouse recombinant M-CSF. BMDM were maintained at 37 °C with 5% CO_2_. On day 4, 10 mL of cDMEM supplemented with 20% LCCM or 40 ng mL^-1^ of M-CSF was added to the BMDM. On day 7, cells were harvested, counted, plated in cDMEM and infected as explained below. When DMEM without glucose was used, the infection was performed after the incubation of BMDM for 2h in glucose-free DMEM.

For ECAR analyses, the bone marrow suspensions were distributed into cryovials at the concentration of 0.5×10^6^ cells mL^-1^ in freezing buffer (90% FBS, 10% dimethyl sulfoxide (DMSO); all GIBCO). For each experiment, cells were thawed and plated at the concentration of 0.5×10^6^ cell/plate in 8 mL of BMDM medium (DMEM medium supplemented with 10% FBS, 5% horse serum, 20% LCCM, 1% HEPES, 1% Sodium Pyruvate and 1% L-glutamine) and differentiated as above.

### Alveolar macrophages isolation

Alveolar macrophages were collected by bronchoalveolar lavage (BAL), performed by exposing the trachea of euthanized mice and injecting 1 mL PBS (GIBCO) + 2% FBS using a 20G catheter connected to a 1 mL syringe. The PBS was flushed into the lung and then aspirated three times and the recovered fluid was placed in a 5 mL tube. The wash was repeated 2 additional times. Cells were plated in complete (c)RPMI (RPMI supplemented with 10% FBS, 1% sodium pyruvate, 1% HEPES and 1% L-glutamine, all GIBCO) supplemented with Penicillin-Streptomycin (100 U mL^-1^; Invitrogen) and allowed to adhere overnight in a 37°C humidified incubator with 5% CO_2_. Media with antibiotics were washed out prior to infection with *M. tuberculosis*.

### Bone-marrow monocytes isolation and cell culture

Bone-marrow monocytes were purified from bone marrow cellular suspensions using an EasySep^TM^ Mouse Monocyte Isolation Kit (STEMCELL Technologies), following the manufacturer’s protocol. Cells were plated in cRPMI supplemented with Penicillin-Streptomycin (100 U mL^-1^; Invitrogen). Media with antibiotics were washed out prior to infection with *M. tuberculosis*.

### In vitro infections

The day before infection, cells were plated at a concentration of 1×10^6^ cell mL^-1^ in 500 μL or 0.5×10^6^ cell mL^-1^ in 200 μL, in 24 or 96 well-plates respectively. Before infection, mycobacterial clumps were disaggregated by gentle passaging through a 25G needle. For BMDM infection, an MOI of 2 bacteria: 1 cell (MOI of 2) was used. Alveolar macrophages and bone marrow monocytes were infected at an MOI of 1. At different time points, cell pellets or culture supernatants were recovered for RNA, protein analysis, or metabolomics. Culture supernatants were filter-sterilized before further processing.

### Chemical inhibitors or agonists

For specific experiments, MCC950 (1 μM; Invivogen), 2-DG (5 mM; Merck), DMM (10 mM; Merck), mitoTEMPO (5 μM; Merck), or rapamycin (10 nM; Merck) was added to BMDM 2h prior infection. Bacteria were treated with 2-DG (5 mM), DMM (10 mM), or Q203 (0.28 nM; MedChemExpress) overnight before infection. The bacteria were then pelleted by centrifugation and the concentration adjusted for infection.

### Cytokine detection

Cytokines were quantified in supernatants of in vitro infected cultures by ELISA (Invitrogen), following the manufacturer’s instructions.

### mRNA analysis by real-time PCR

Total RNA from infected cells was extracted with TripleXtractor (GriSP Research Solutions), according to the manufacturer’s instructions and quantified using the NanoDrop 1000 Spectrophotometer (Thermo Scientific). Complementary deoxyribonucleic acid (cDNA) was synthesized using the ProtoScript First-Strand cDNA Synthesis kit (New England Biolabs). Real-time PCR assays were performed using a CFX Connect Real-Time PCR Detection System (Bio-Rad). Target gene mRNA expression was quantified by real-time PCR using iTaq Universal SYBR Green Supermix (Applied Biosystems) and normalized to ubiquitin mRNA levels. The oligonucleotide sequences used are listed in S1 Table. The specificity of amplification was confirmed by melting curve analysis.

### Lactate quantification

Culture supernatants were collected 24h post-infection for the measurement of lactate release, using the Lactate Assay Kit (Merck) accordingly to the manufacturer’s instructions.

### Cell viability quantification

24h post-infection, Orangu^TM^ (Cell Guidance Systems) was added to the cells and incubated for 1-3h at 37 °C, accordingly to the manufacturer’s instructions. Absorbance was read at 450 nm.

### High-performance liquid chromatography (HPLC)

Glucose and lactate were quantified in cell culture supernatants by HPLC (Gilson bomb system); HyperREZ XP Carbohydrate H+ 8μM as before (49). Briefly, samples were filtered with a 0.2 μm filter, the mobile phase (0.0025M H_2_SO_4_) refiltered and degasified for 30 min, and analysed using the following running protocol (sensitivity 8): 15 min at a constant flux of 0.7 mL/min at 54°C for glucose and lactate detection. The peaks were detected in a refractive index detector (IOTA 2, Reagents) and integration was performed using Gilson Uniprot Software, version 5.11.

### Extracellular flux analysis

Differentiated BMDM were plated in XFp plates (Agilent), previously coated with Cell-Tak (Corning), at the concentration of 1×10^6^ cells mL^-1^ in 80 μL of BMDM medium. Plates were incubated at 37 °C with 5% CO_2_ overnight before infection. Cell-Tak was applied to the plates according to the manufacturer’s protocol. For the infection, stocks of *M. tuberculosis* 4I2 or *M. tuberculosis* 6C4 were thawed, resuspended and added to Middlebrook 7H9 liquid medium complemented with 10% OADC, 2% glycerol. The culture was incubated at 37 °C with constant 120 rpm shaking until mid-log phase. For these experiments, the cultures used were in passages 2 to 5. Culture optical density (OD) was measured a day before infection and concentration was adjusted. On the day of infection, bacteria were washed, resuspended with a syringe with a sub-Q needle, and the concentration adjusted in cDMEM.

ECAR was measured using the Seahorse XFp extracellular flux analyser (Agilent). The assay performed was the glycolysis stress test. The Wave Desktop 2.6.0 Software was used to normalize the ECAR values to the cell number of each well and export the data into GraphPad Prism Version 7.04 for statistical analysis and calculation of the parameters (glycolysis, glycolytic capacity and glycolytic reserve). Basal ECAR readings were obtained. After the injection of each compound, ECAR readings were also obtained. The following compounds were added to the injection ports: glucose 10 mM (Agilent), oligomycin 2 μM (Agilent), 2-DG 50 mM (Sigma). Growth media was exchanged with XF DMEM medium, without phenol red (Agilent), supplemented with 2 mM L-glutamine, 1h before the assay. The Agilent Seahorse Sensor Cartridge was hydrated one day before the assay and calibrated right before the assay, according to the manufacturer’s protocol. Assays were normalized through total cell numbers. After the assay, cells were fixed overnight In 4% PFA. Cells were then stained with 4,6-diamidino-2-phenylindole (DAPI). Images were acquired on the Cell Insight CX7 High-Content Screening (HCS) and subsequently analysed using an in-built software, HCS Studio (ThermoFisher Scientific).

### Mitochondrial membrane potential quantification

The CMXRos dye was prepared as per the manufacturer’s recommendation (Invitrogen), dissolved in DMSO at a concentration of 1 mM and stored at -20 °C until use. Cells were incubated at 37 °C with 5 μM of CMXRos for 30 min in the dark, 24h post-infection. Cells were then harvested, fixed in 4% PFA for 20 min, and washed. Samples were acquired in the Accuri C6 Plus. MFI were analysed using FlowJo (version 10.1.r7).

### THP1-ASC-GFP differentiation, infection and speck quantification

The THP1-ASC-GFP (InvivoGen) cell line was maintained in cRPMI supplemented with 100 μg mL^−1^ Zeocin (InvivoGen) and 100 U mL^-1^ Penicillin-Streptomycin (Invitrogen), at 37 °C and 5% CO_2_. Cells were used before reaching 20 passages. THP1-ASC-GFP cells were differentiated with 100 nM PMA for 24h and allowed to rest for 4 days in fresh medium without PMA, before being infected at an MOI of 1. For immunofluorescent detection of ASC specks, infected cells were washed with PBS and fixed in 4% formalin. Fixed cells were stained with PhenoVue Hoechst 33342 (NOVA Natura) 10 μg mL^-1^ and HCS CellMask Deep Red (Invitrogen) 10 μg mL^-1^. Plate image acquisition was performed with an Operetta CLS High-Content Analysis System, with the objective 20x Water, NA 1.0. Images were analysed using Harmony software.

### NMR-based metabolomics

Supernatants were collected 24h post-infection and stored at −80°C; cells were washed 3 times with cold PBS and immediately extracted using a biphasic extraction protocol with methanol/chloroform/water (1:1:0.7), as previously described (50). The resulting polar extracts were dried under vacuum in a SpeedVac concentrator (Eppendorf) and stored at −80°C until analysis. Cell-conditioned media was thawed and subjected to a protein-precipitation procedure using cold methanol (51). The resulting supernatants were vacuum-dried and stored at −80°C until NMR analysis. The respective cell-free medium was incubated and processed in the same conditions and used to assess metabolite consumption and excretion by cells. For NMR analysis, the dried samples were resuspended in 600 mL of deuterated PBS (100 mM, pH 7.4) containing 0.1 mM 3-(trimethylsilyl) propanoic acid (TSP-*d_4_*), and 550 mL of each sample were transferred to 5 mm NMR tubes. All samples were analyzed in a Bruker Avance III HD 500 NMR spectrometer (University of Aveiro, Portuguese NMR Network), operating at 500.13 MHz for ^1^H observation, at 298 K, using a 5 mm TXI probe. Standard 1D ^1^H spectra with water presaturation (pulse program “noesypr1d” from Bruker library) were acquired and processed as previously described (52). Metabolite assignment was based on matching spectral information to reference spectra available in Chenomx (Edmonton), BBIOREFCODE-2–0–0 (Bruker Biospin) and the Human Metabolome Database (HMDB) (53). Principal Component Analysis (PCA) was carried out in MetaboAnalyst 5.0.

### In silico study of *M. tuberculosis* SDH-1

Amino acid sequences used to predict the 3D structure of *M. tuberculosis* SDH-1 were obtained from the NCBI protein database. Sequences for *M. tuberculosis* H37Rv were the Flavoprotein subunit (NP_214762.1), Iron-Sulphur subunit (NP_217836.1), and Membrane anchor subunit (NP_214763.1). Sequences were exported to ColabFold (https://colab.research.google.com/github/sokrypton/ColabFold/blob/main/AlphaFold2.ipynb#scrollTo=kOblAo-xetgx) (39). Models for each domain were individually built against pdb100 templates, and the five best models were relaxed using AMBER software included in ColabFold. The best models were analysed, and superpositioned with *M. smegmatis* SDH-1 (PDB: 7D6V) using the PyMOL program (The PyMOL Molecular Graphics System, Version 2.5.2, Schrödinger, LLC). Small molecule structures for Ubiquinone-1 (CID 4462), DMM (CID 7943), Malonate (CID 9084) and Succinate (CID 160419) were extracted from PubChem.

### Statistical analysis

Results are represented as mean ± standard deviation (SD) or in box and whiskers plots (Tukey’s method). Data were analysed using GraphPad Prism software, version 9.0. Reference to “n” is in the Figure legends. Data was checked for normality and log normality with Shapiro-Wilk or D’Agostino-Pearson tests, as well as for outliers with Grubbs’s test. Based on this information, parametric or nonparametric statistical tests were used for multiple corrections and correlations. Differences were considered significant for p≤0.05 and indicated in the figures.

## Acknowledgements

We thank Drs Gil Castro, Tiago Beites and Joana Couto for critically reading the manuscript. The authors thank the support from the i3s scientific platforms animal facility; translational cytometry and biosciences screening, a member of the PT-OPENSCREEN (NORTE-01-0145-FEDER-085468) and PPBI (PPBI-POCI-01-0145-FEDER-022122).

This work was funded by the FCT grant PTDC/SAU-INF/1172/2021 and CEECIND/00241/2017 to MS. AIF was funded by the Portuguese Foundation for Science and Technology (FCT) PhD scholarship 2020.05949.BD, an EFIS-IL Short-term fellowship and a Research Grant 2022 from the European Society of Clinical Microbiology and Infectious Diseases (ESCMID). DS held a FCT PhD fellowship with reference SFRH/BD/143536/2019. This work was also developed within the scope of the project CICECO – Aveiro Institute of Materials UIDB/50011/2020 (DOI 10.54499/UIDB/50011/2020), UIDP/50011/2020 (DOI 10.54499/UIDP/50011/2020) & LA/P/0006/2020 (DOI 10.54499/LA/P/0006/2020), financed by national funds through the FCT/MEC (PIDDAC), and projects 10.54499/UIDB/50026/2020, 10.54499/UIDP/50026/2020 and 10.54499/LA/P/0050/2020. The NMR spectrometer is part of the National NMR Network (PTNMR), partially supported by Infrastructure Project N° 022161 (co-financed by FEDER through COMPETE 2020, POCI, and PORL and FCT through PIDDAC). IFD and RS thank FCT for the contracts CEECIND/02387/2018 and 10.54499/2020.00185.CEECIND/CP1600/CT0004, respectively.

## Supplementary Figure Legends

**S1 Fig.** C57BL/6 BMDM were pre-treated for 2h with MCC950 (**A, B, F**), 2-DG (**C, F**), DMEM without glucose (**C, F**), or 6AN (**E, F**) and subsequently infected with either *M. tuberculosis* clinical isolate. (**A**) 6h and 15h post-infection, cell cultures were lysed and the expression of *Il1b* measured by real-time PCR. The amount of **(B, E)** secreted IL-1β or (**C**) lactate was quantified in the cell culture supernatants 24h post-infection. (**D**) 6AN inhibitory action within the pentose phosphate pathway. (**F**) Cell viability was measured by ORANGU cell counting solution colorimetric assay. Data are represented in log_2_(fold change), with 0 representing macrophages infected with *M. tuberculosis* 4I2 or -6C4. Represented is the mean ± SD (**A-C, E**) or box and whiskers plots (Tukey’s method) (**F**) of triplicate (**A-F**) wells for two experiments (**E, F**) or one experiment representative of two (**A-C**). Statistical analysis was performed using a two-tailed unpaired Student’s *t*-test. 2-DG – 2-deoxy-glucose, 6AN – 6-aminonicotinamide, G6PD – glucose-6-phosphate dehydrogenase, Mtb – *M. tuberculosis*, NI – non-infected.

**S2 Fig.** C57BL/6 BMDM were infected with either clinical isolate at an MOI of 2. (**A**) 24h post-infection, BMDM were stained with Mitotracker Red CMXRos and MFI were analysed and are represented in the figure. (**B**) 6h and 15 h post-infection, cell cultures were lysed, and the expression of the indicated genes analysed by real-time PCR. (**C**) Cell viability for BMDM pre-treated for 2h with the indicated inhibitors measured 24h post-infection using a colorimetric assay. Data are represented in log_2_(fold change), with 0 representing macrophages infected with *M. tuberculosis* 4I2 (right panel) or 6C4 (left panel). (**D**) THP1-ASC-GFP macrophages were infected with *M*. tuberculosis 4I2 alone, after 2h of pre-treatment with MCC950 or with *M. tuberculosis* 6C4 alone at an MOI of 1 for 6h. 60 fields (total number of cells indicated in each bar) per replicate were acquired and analysed. The percentage of speck-positive cells was determined for the different conditions. Represented is the mean ± SD (**B, D**) or box and whiskers plots (Tukey’s method) (**C**) of triplicate wells for one experiment. Statistical analysis was performed using two-tailed unpaired Student’s *t*-test (**B**) or One-way ANOVA (**D**). DMM – dimethyl malonate, Mtb – *M. tuberculosis*, NI – non-infected, p.i. – post-infection.

**S3 Fig.** C57BL/6 BMDM were infected with *M. tuberculosis* 4I2 or 6C4 or left uninfected and 24h later the supernatant and cellular extract recovered for NMR metabolomics. In parallel, *M. tuberculosis* 4I2 and 6C4 cultures were also recovered. (**A**) Metabolites consumed and excreted among the tested conditions. Variations are expressed in relation to acellular media. (**B**) Relative intracellular levels of metabolites. Represented are box and whiskers plots (Tukey’s method) of duplicate wells for two experiments. Each replicate is a pool of 24 wells in a 24 well-plate. Statistical analysis was performed using One-way ANOVA. au – arbitrary unit, BMDM – bone-marrow derived macrophages, Mtb - *M. tuberculosis*, m – medium, NI – non-infected.

**S4 Fig.** (**A**) *M. tuberculosis* clinical isolates or BMDM from C57BL/6 mice were treated with DMM prior to infection according to the protocol in **Fig 5B**. Treated macrophages were infected with non-treated bacteria and non-treated macrophages were infected with treated bacteria, at an MOI of 2 for 6h and 15h. At the indicated timepoints, cell cultures were lysed and the expression of *Il1b* measured by real-time PCR. Represented is the mean ± SD of triplicate wells for one experiment. Statistical analysis was performed using a two-tailed unpaired Student’s *t*-test. BMDM – bone-marrow derived macrophage, Mtb – *M. tuberculosis*, DMM – dimethyl malonate.

**S1 Table.**
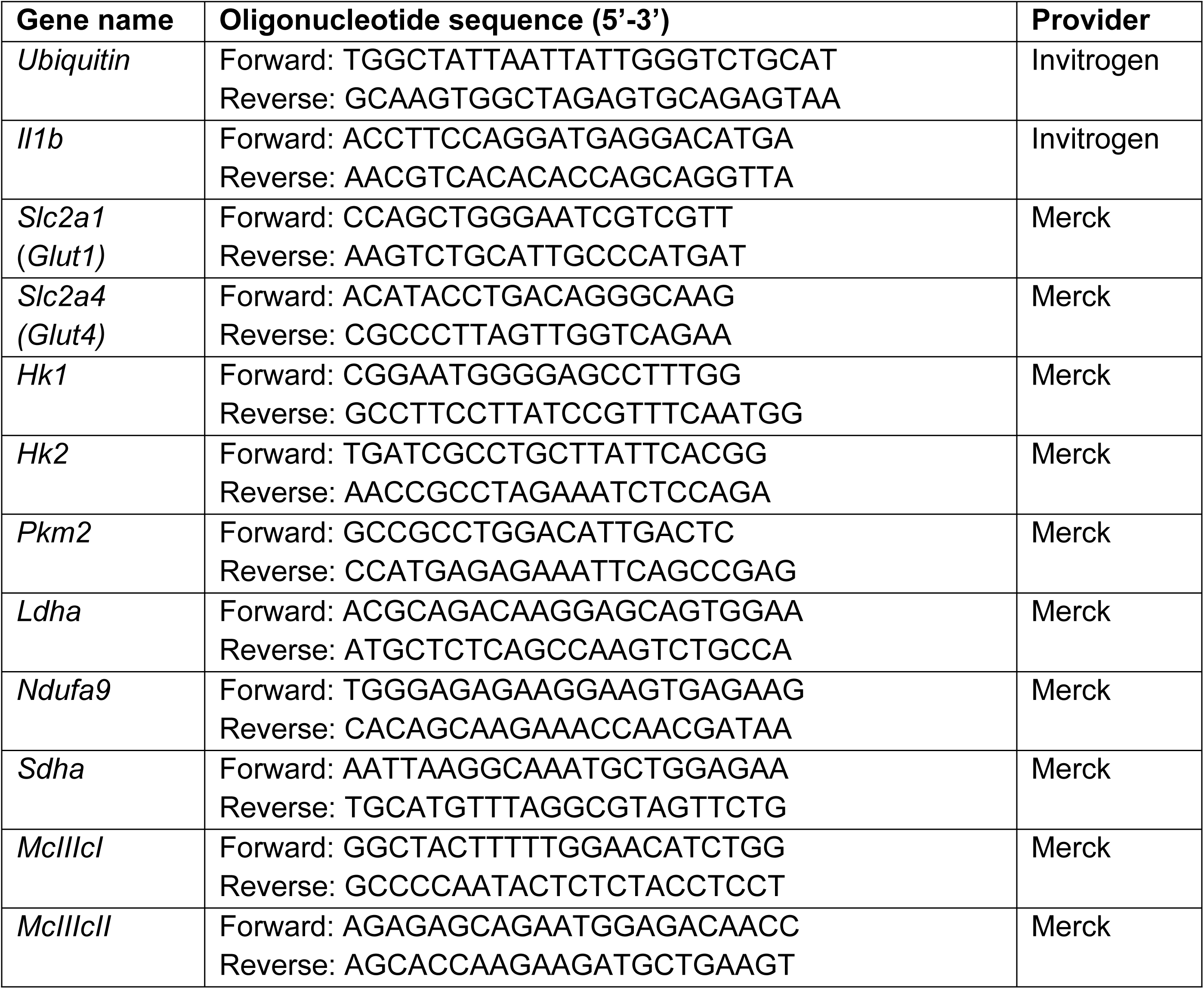
List of oligonucleotide sequences.

